# Sequential experience reshapes population representations in visual cortex

**DOI:** 10.64898/2026.03.18.712658

**Authors:** Lily E. Kramer, Marlene R. Cohen

## Abstract

Visual experience is organized in time. When riding the same bus route each day, the visual scene unfolds in a predictable order without requiring active choice. During goal-directed behavior, individuals organize actions into routines, such as repeatedly walking the same route to work even when alternatives are equally efficient. Because experience unfolds across sequences of events, identifying how it reshapes population activity requires examining representations over time. Many studies have shown that repeated experience reduces mean firing rates in visual cortex^1–14^. While firing rates effectively signal novelty or repetition, they are not well positioned to describe how populations of neurons represent temporal relationships. A growing body of work suggests that the geometry of population activity provides additional insight into how visual information is structured and read out^15–26^. We examined how experience with temporal structure reshapes the geometry of population activity in visual area V4. We recorded neuronal populations across three contexts that varied in temporal structure and behavioral relevance: repeated presentation of individual images, passive exposure to structured image sequences, and repeated execution of self-chosen visually guided action sequences for reward. Across contexts, experience constrained population responses toward a typical activity pattern. In sequence contexts, experience made temporal position more linearly accessible and, during active practice, increased the separability of task-relevant variables. These findings show that experience reorganizes the geometry of visual population activity to reflect temporal structure, constraining responses and altering how sequence-related information is represented.

## Introduction

Experience-dependent changes in visual cortex are typically studied using repeated presentation of individual images. Across many visual areas, responses to novel or unexpected stimuli are enhanced, whereas repeated stimuli elicit reduced responses, a phenomenon commonly described as repetition suppression^1–14^. These effects are robust and have been documented in many studies and visual areas.

Although repetition suppression demonstrates that experience modulates visual neuronal responses, visual experience in natural settings extends beyond repetition of isolated stimuli. Visual input often unfolds in ordered sequences, and behavior frequently involves repeated action routines. In such temporally structured contexts, experience may shape not only response magnitude but also how information about order, sequence position, and task context is organized across populations of neurons.

Analyses focused on firing rate typically consider neurons individually and emphasize changes in mean response magnitude. While such measures reveal reliable effects of familiarity, they do not characterize how population activity encodes relationships among events over time. Recent work suggests that the geometry of population activity provides additional insight into how information is encoded and read out^15–19^. Such analyses quantify how responses are organized relative to their mean activity pattern, for example using multivariate measures such as Mahalanobis distance^20,21^ or dimensionality^21–23^ and how different task-relevant variables are represented within the population^24–26^. These approaches provide a framework for examining how experience reshapes representations beyond changes in mean firing rate.

To test this idea, we examined how experience with temporal structure is associated with changes in population activity in visual area V4, a mid-level visual area that encodes diverse visual features and is modulated by attention, task demands, and learning^3,15,17,27–36^. We studied three contexts that varied in temporal structure and behavioral relevance: image familiarity, passive exposure to structured image sequences, and repeated execution of visually guided action sequences for reward. These paradigms allowed us to assess whether experience in each setting is accompanied by systematic changes in the geometry of population activity.

In a classic familiarity paradigm, we replicated prior observations that novel stimuli elicit higher firing rates and extended this analysis to the population level by quantifying how responses are organized relative to their mean activity pattern. Across all three contexts, increasing experience was associated with a reduction in the distance of population responses from the mean, measured using Mahalanobis distance. In the sequence contexts, we further asked whether this reorganization reflected learned temporal relationships. In both passive and action-based sequence exposure, population activity reorganized so that temporal position became more linearly accessible. In the action sequence task, which included reward contingencies and multiple task-relevant variables, we additionally tested whether experience altered how these variables were represented and found that population activity became more separable across task-relevant dimensions. Together, these experiments document experience-dependent changes in the geometry of visual population activity that relate to learned temporal structure.

## Results

### Passive image experience reduces and constrains population responses in area V4

We began by examining passive image familiarity, a well-established paradigm for characterizing how visual experience influences responses in visual cortex. We presented grayscale images of objects and animals to two monkeys while recording from neuronal populations in mid-level visual area V4 using chronically implanted multielectrode arrays. We chose V4 because it is a primarily visual area whose responses are modulated by cognitive processes including different forms of learning^15,29,31,32^, and because prior work has shown that V4 responses are modulated by image familiarity^3^. On each trial, we presented a single image within the joint receptive fields of the recorded units (Figure 1a). Each image was presented on two subsequent trials. To minimize low-level adaptation, we rewarded the monkey for making an eye movement to a subsequently presented target at a random location that did not overlap the image after stimulus offset.

**Figure 1.**
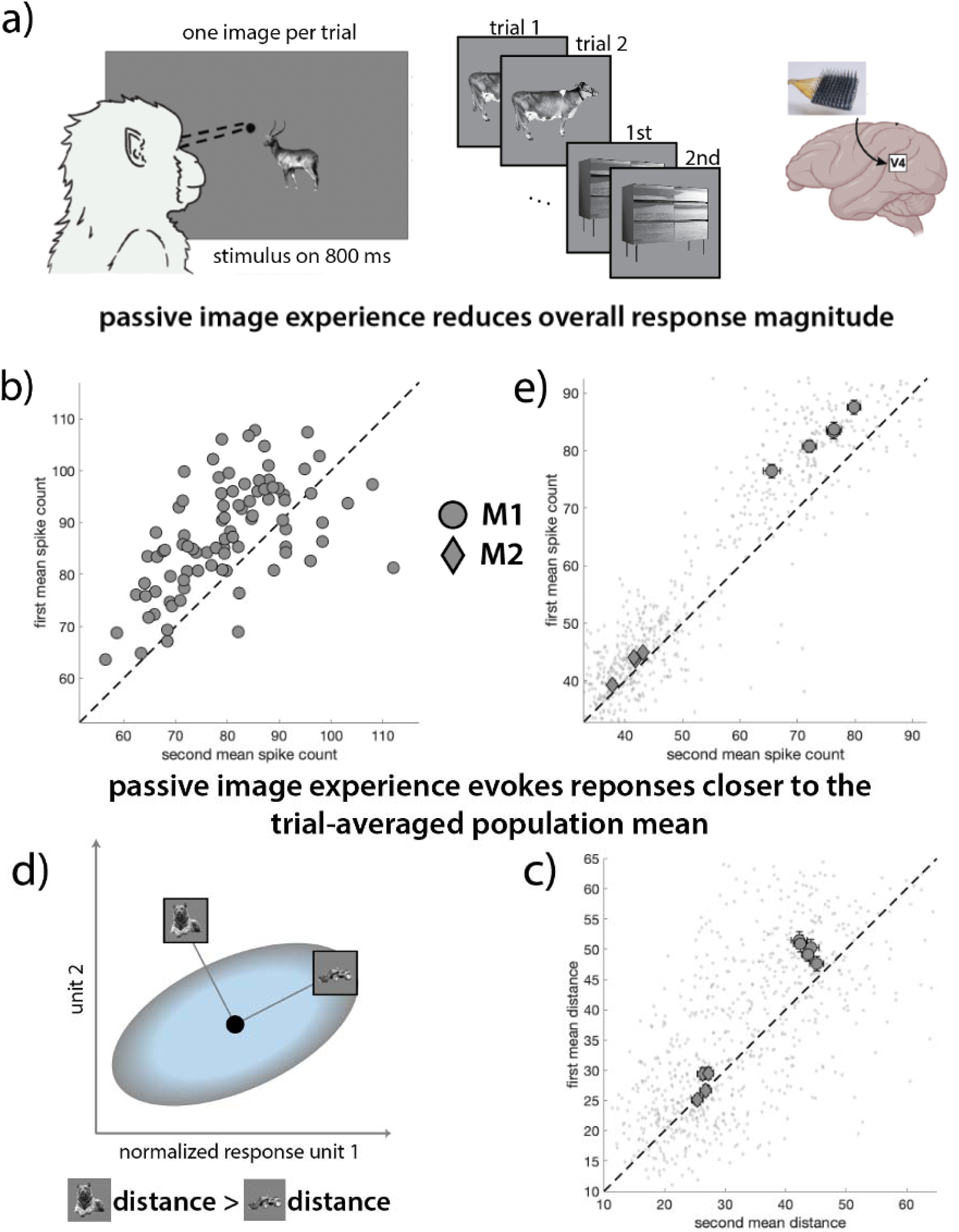
Passive image experience experiment: **a) Task and electrophysiology.** On each trial, animals passively viewed an image presented in the joint receptive fields of the recorded V4 units while fixating a central point. Following the stimulus presentation period (860ms), animals were rewarded for making a saccade to a random, radial location that did not overlap the image. We analyzed responses during back-to-back presentations of the same image. **b) Response reduction for each image in a single example session (monkey M1)**. Each point is the mean population spike count in response to the first (y-axis) and second (x-axis) presentations of a distinct image. Points above the unity line indicate reduced responses on the second presentation. **c) Average response reduction across sessions.** Circular points represent mean responses (averaged over images and neurons) for each session for monkey M1, and diamonds are from monkey M2. Error bars represent standard error of the mean (SEM). **d) Schematic illustrating the Mahalanobis distance metric.** In a simplified two-dimensional population space (where each axis is the normalized response of one V4 unit), the Mahalanobis distance is the distance between the population response to each image and the trial-averaged population response (central, black point), normalized by the covariance matrix (ellipse). This normalization means that points that are more out of distribution (e.g. responses to the tiger) have higher Mahalanobis distance than within-distribution responses (e.g. to the tractor). **e) Average Mahalanobis distance on first (y-axis) and second (x-axis) presentations for each session, averaged over images.** Points above the unity line indicate more constrained population responses on repeated presentations of an image. Conventions as in (b).

Consistent with previous work^3^, repeated image presentations evoked weaker average responses (the mean spike counts for most sessions fall above the unity line in Figure 1b,c, indicating stronger responses to first compared to second presentations of each image). This effect was robust in both animals (one-sided, paired t-tests: M1 p = 1.7×10^−48^, effect=0.96; M2 p = 1.1×10^−8^, effect=0.28).

To examine how passive image experience affects population organization, we quantified the Mahalanobis distance between the population response on each trial and the trial-averaged response across all stimuli. This measure captures how responses are distributed relative to their mean activity pattern while accounting for covariance between neurons (Figure 1d). Repeated image presentations evoked population responses that were closer to the mean than first presentations (Figure 1e). This reduction in distance was significant in both animals (one-sided, paired t-test: M1 p = 1.0×10^−16^, effect = 0.48; M2 p = 0.012, effect = 0.11). Changes in spike count and Mahalanobis distance were moderately correlated across images (M1 r = 0.41; M2 r = 0.34), indicating that reductions in response magnitude were accompanied by systematic changes in population organization.

Together, these results show that passive image experience is associated not only with reduced response magnitude but also with a constraint of population activity toward a more typical activity pattern. In this paradigm, familiarity is associated with a compression of population responses in population space. However, because the images were not organized into a structured sequence and no task depended on their order, this paradigm does not reveal whether such geometric changes reflect learned temporal relationships or have functional consequences for behavior. We therefore turned to additional experiments involving structured sequences to examine how experience-dependent changes in population geometry relate to temporal organization.

### Passive sequence experience reorganizes population responses to reflect learned temporal order

One function of learning is to enable prediction: when sensory events occur in a reliable temporal order, the brain can anticipate upcoming stimuli and prepare appropriate responses^37–42^. In primates, previous studies have shown that serial-order associations affect neuronal responses of higher-level visual processing areas^8,10,11,43–45^. We hypothesized that experience with temporal structure should reshape mid-level sensory representations to reflect expected sequence position, beyond familiarity with the images themselves. To test this idea, we examined how learning the temporal order of already familiar images affects population responses in V4.

We presented sequences of images to three monkeys while recording neuronal population activity in visual area V4. On each trial, animals were rewarded for maintaining central fixation while one image sequence was presented. All images appeared at the same location, overlapping the joint receptive fields of the recorded V4 units (Figure 2a). Two sequences of four images were constructed either from grayscale images of animals and objects (Figure 2a, top row) or colored, curved 3D shapes (Figure 2a, bottom row). These same sequences were reused across sessions, allowing animals to become familiar with their temporal order.

**Figure 2.**
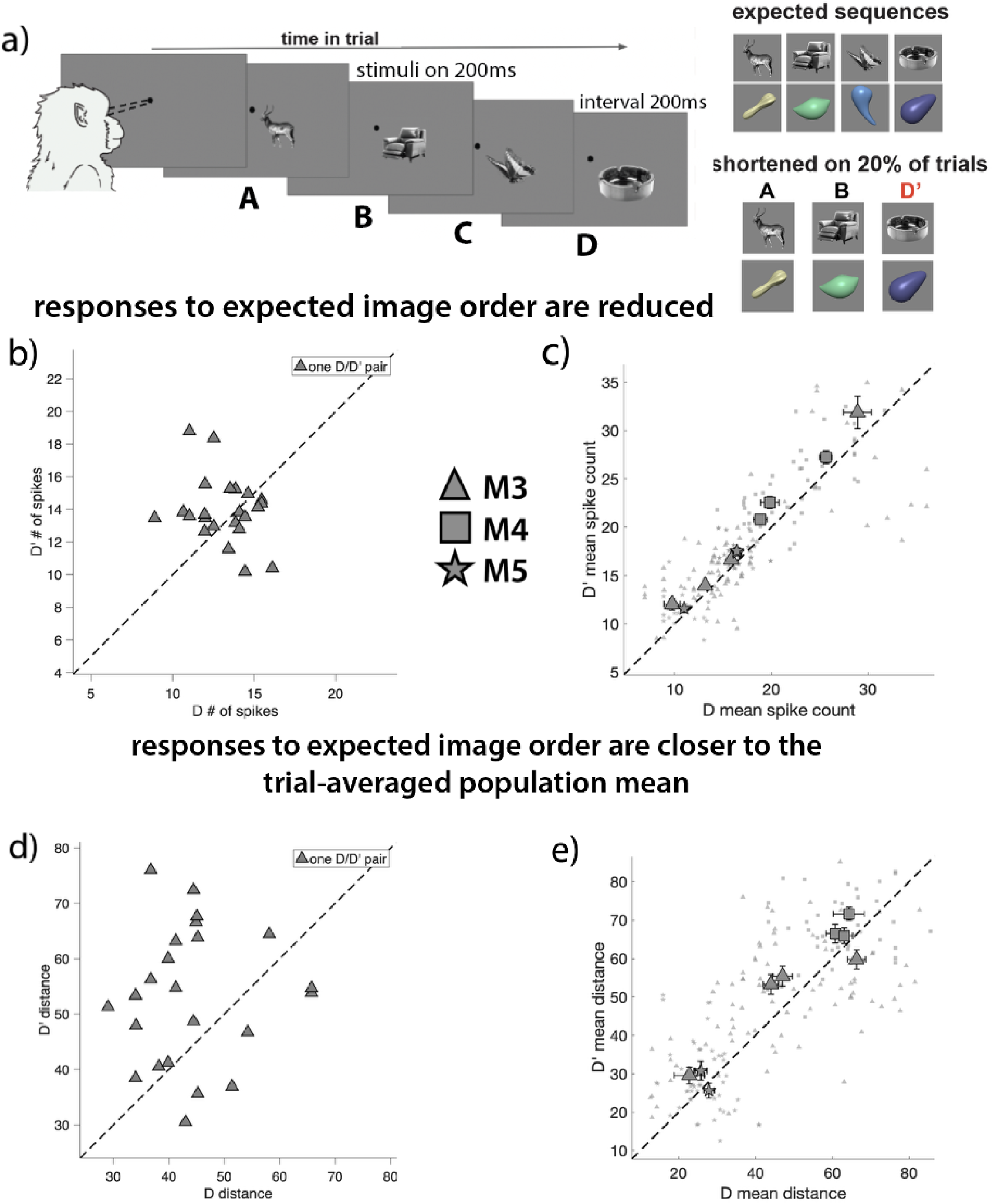
Passive visual-temporal association experiment: **a) Task and electrophysiology.** On each trial while fixating on a central point, animals passively viewed one sequence of images (A, B, C, D) presented at the location of their V4 unit receptive fields. Each image in the sequence was presented for 200ms, with a 200ms interval between images. At the end of each trial (sequence), the animal was rewarded for fixating. On a randomly selected 20% of trials, the third stimulus (C) was omitted and the fourth stimulus (D’) was presented in its place. The same two expected sequences (one for each stimulus set) were used in all sessions. **b) Average population responses to the same image when it occurred in the unexpected (D’; y-axis) or expected (D; x-axis) sequence position in a single example session (monkey M3).** Each point is the unit-averaged spike count response to one presentation of D’ in comparison to the closest preceding presentation of D. **c) Average (over exposure pairs) population spike counts for each session.** Error bars represent the SEM. **d) Mahalanobis distances for expected and unexpected D presentations in a single example session (monkey M3).** Each data point is the distance from the trial-averaged, mean population response to all D’ (y-axis; unexpected third position in the sequence) and D (x-axis; expected, fourth position in the sequence). **e) Average (over exposure pairs) Mahalanobis distances for each session.** Conventions as in (c).

Within each session, the expected sequence was presented on approximately 80% of trials. On the remaining 20% of trials, a sequence violation occurred: the third image (C) was omitted, and the fourth image (D’) appeared in the third position (Figure 2b). This manipulation allowed us to assess whether V4 responses were sensitive to learned temporal associations despite passive viewing.

We compared responses to the final image (D or D’, referring to whether it appeared in the fourth or third position, respectively; see Methods for pairing procedure). Responses to the expected image D were reduced relative to unexpected presentations D’ (Figure 2b, c), and this reduction was significant in all three animals (one-sided paired t-tests: M3 p = 0.020, effect = 0.17; M4 p = 1.3×10^−4^, effect = 0.38; M5 p = 0.0075, effect = 0.22). As a control, responses to the second image (B) were indistinguishable on violation and non-violation trials in all animals (one-sided paired t-tests: M3 p = 0.78; M4 p = 0.63; M5 p = 0.70). This is expected because the animals could not anticipate violations.

To assess whether temporal position influenced population geometry, we compared Mahalanobis distance for D and D’. Presentations at the expected position (D) were closer to the trial-averaged population mean response than unexpected presentations (D’) (Figure 2d, e). This difference was significant in two animals (M3 p = 0.028, effect = 0.20; M4 p = 0.0025, effect=0.50) and approached significance in the third (M5 p=0.091, effect=0.20). No significant differences were observed for B across violation conditions (M3 p = 0.18; M4 p = 0.15; M5 p = 0.99).

These results demonstrate that passive experience with temporal structure modulates V4 population responses even when image familiarity is held constant. Images occurring at expected sequence positions evoke lower firing rates and more constrained population representations than the same images occurring at unexpected positions. Thus, learning temporal order reorganizes visual population responses in ways that cannot be explained by familiarity with the image itself.

### Passive sequence experience organizes V4 population geometry to better and more linearly encode temporal order

The position-dependent modulation observed above indicates that experience with temporal order influences V4 responses beyond image familiarity. However, changes in response magnitude and Mahalanobis distance do not by themselves reveal how sequence order is represented within population activity. Theoretical work suggests that learning temporal structure supports prediction^37,39–41,46–50^. If so, experience should organize population activity such that information about sequence position is embedded in population geometry, rather than reflected solely as changes in firing rate.

Before examining sequence structure, we confirmed that passive exposure produced the expected reduction in mean response over the course of each session. Averaging population responses across all four sequence elements (A–D), responses during late trials were reduced relative to early trials (Figure 3a). This reduction was significant in two of three animals (M3 p = 0.13, effect = 0.84; M4 p = 0.0015, effect = 2.0; M5 p =2.3×10^−5^, effect = 1.3).

**Figure 3.**
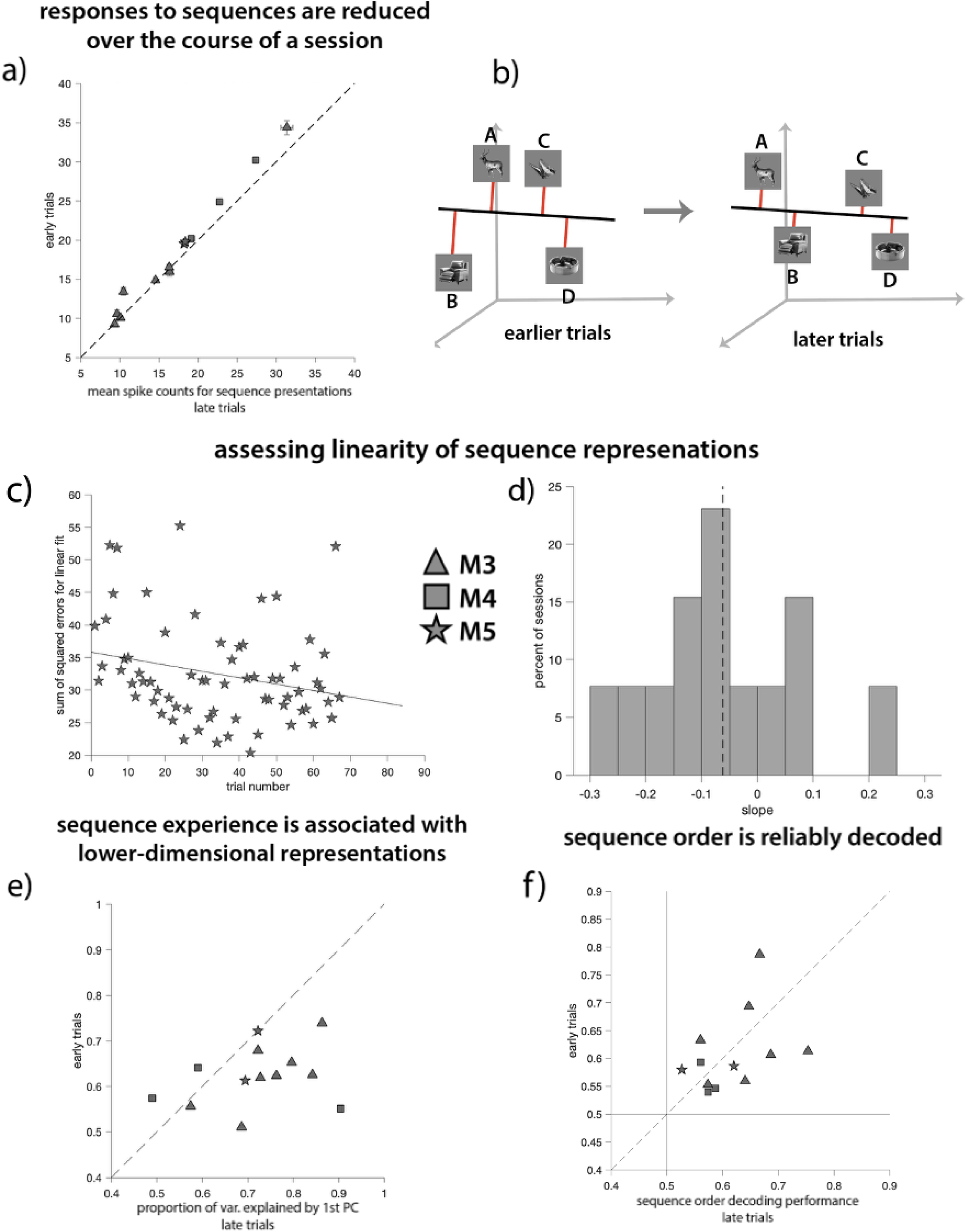
Passive experience with visual-temporal associations organizes population activity according to sequence order: **a) Population responses decrease over the course of a session.** Each point shows the mean population response for one session, averaged over neurons and over all four images in the expected sequence (A-D), comparing the early (first 25 trials) and late (next 25 trials) **sequence presentations.** Error bars represent the SEM. **b) Schematic illustrating experience-dependent organization of population responses.** Mean population responses to the four sequence elements define an ordered trajectory in population space (black line). With experience, single-trial responses become better aligned with this trajectory, reflected in smaller residuals (red lines) and a lower dimensional representation. **c) Example session (M5) showing the sum of squared residuals for each sequence presentation relative to the line fit to the mean population responses for A–D.** Each point corresponds to one trial; decreasing residuals over trials indicate increasing alignment with the sequence axis. **d) Distribution of slopes describing changes in residual magnitude over trials across sessions.** Slopes are obtained by fitting a line to the residuals in panel (c) for each session. Negative slopes indicate increasing alignment of population responses with the sequence axis over time. The dashed line denotes the mean slope across sessions. **e)Experience-dependent reduction in response dimensionality.** Each point shows the proportion of variance explained by the first principal component for early (y-axis) and late (x-axis) trials within a session. Values below the unity line indicate increased variance explained late in the session. **f) Sequence order decoding performance for each session using a linear classifier trained and tested on either early or late trials.** Performance above 0.5 indicates above-chance decoding; the unity line compares early and late trials within each session.

We then tested whether population responses became more structured with respect to temporal order using three complementary analyses.

First, we asked whether responses to the four sequence elements (A–D) aligned along a single axis in population space and whether this alignment strengthened over time. For each session, we fit a line to the mean population responses for A–D and computed residuals for single-trial responses relative to this axis. If experience organizes activity according to sequence order, residuals should decrease across trials. Across sessions, slopes describing residual magnitude tended to be negative (mean = –0.0619, SEM = 0.0360), consistent with increasing alignment to a sequence axis, though this effect did not reach significance (one-sided two-sample t-test: p = 0.056; Figure 3c, d).

Second, we examined whether the dimensionality of population responses changed with experience. We calculated the proportion of variance explained by the first principal component for early and late trials. Across sessions, late trials exhibited greater variance explained by the first component (Figure 3e), indicating reduced dimensionality with experience (one-sided paired t-test: p = 0.0054, effect = 0.098).

Third, we assessed whether population activity reflected sequence order in a structured manner beyond dimensionality reduction. Using a cross-condition generalization analysis, we trained linear classifiers on one partition of the sequence and tested whether the learned decision boundary generalized to held-out elements in a manner consistent with sequence order. High generalization across complementary train-test splits indicates that a single projection of population activity supports multiple order-consistent distinctions.

Importantly, this analysis dissociates low dimensionality from ordered structure: responses may vary along one dominant dimension yet fail this test if sequence elements are arranged non-monotonically along that dimension. Sequence order was decoded reliably above chance from both early and late trials (Figure 3f; early p = 8.2×10^−5^, effect = 0.10; late p = 1.7×10^−5^, effect = 0.11). The early–late difference was not significant (p = 0.33).

Together, these analyses show that passive experience organizes V4 population activity such that temporal relationships among stimuli are embedded in population geometry, even in the absence of active task demands.

### Learning action sequences links temporal structure to behavior

Our passive sequence experiments demonstrate that temporal order is reflected in population activity in visual cortex, even in the absence of explicit task demands. However, because those experiments required no action selection and had no behavioral consequences tied to sequence structure, they do not establish whether experience-dependent changes in population activity are related to improvements in behavior. To address this limitation, we designed a task in which successful performance depends directly on learning and executing a sequence of visually guided actions, allowing us to examine how experience with structured action sequences shapes both behavior and visual population responses.

We trained an adult rhesus macaque (monkey M3 from the passive image sequence experiment) to perform a task requiring the active formulation and practice of sequences of actions (eye movements), while recording from populations of V4 units using the same recording methods as in previous experiments (see Methods).

Prior to the analyzed sessions, the animal was trained on a simplified version of the task to learn its structure and rules.

On each trial, following an initial fixation period, the animal was presented with a grid of twenty-five targets arranged in a two-dimensional array (Figure 4a). A game piece appeared at one grid location, and a rewarded target appeared at another. Fixation requirements were then relaxed, and the animal was free to move the game piece toward the rewarded stimulus using sequences of eye movements. Each move required fixation on the game piece (200 ms), a saccade to one of four adjacent targets, and maintained fixation at the selected location (200 ms). Correct moves updated the game piece location. The trial terminated with juice reward when the game piece reached the rewarded location, or without reward if twenty seconds elapsed. The animal could revisit previously occupied locations, allowing flexible exploration.

**Figure 4.**
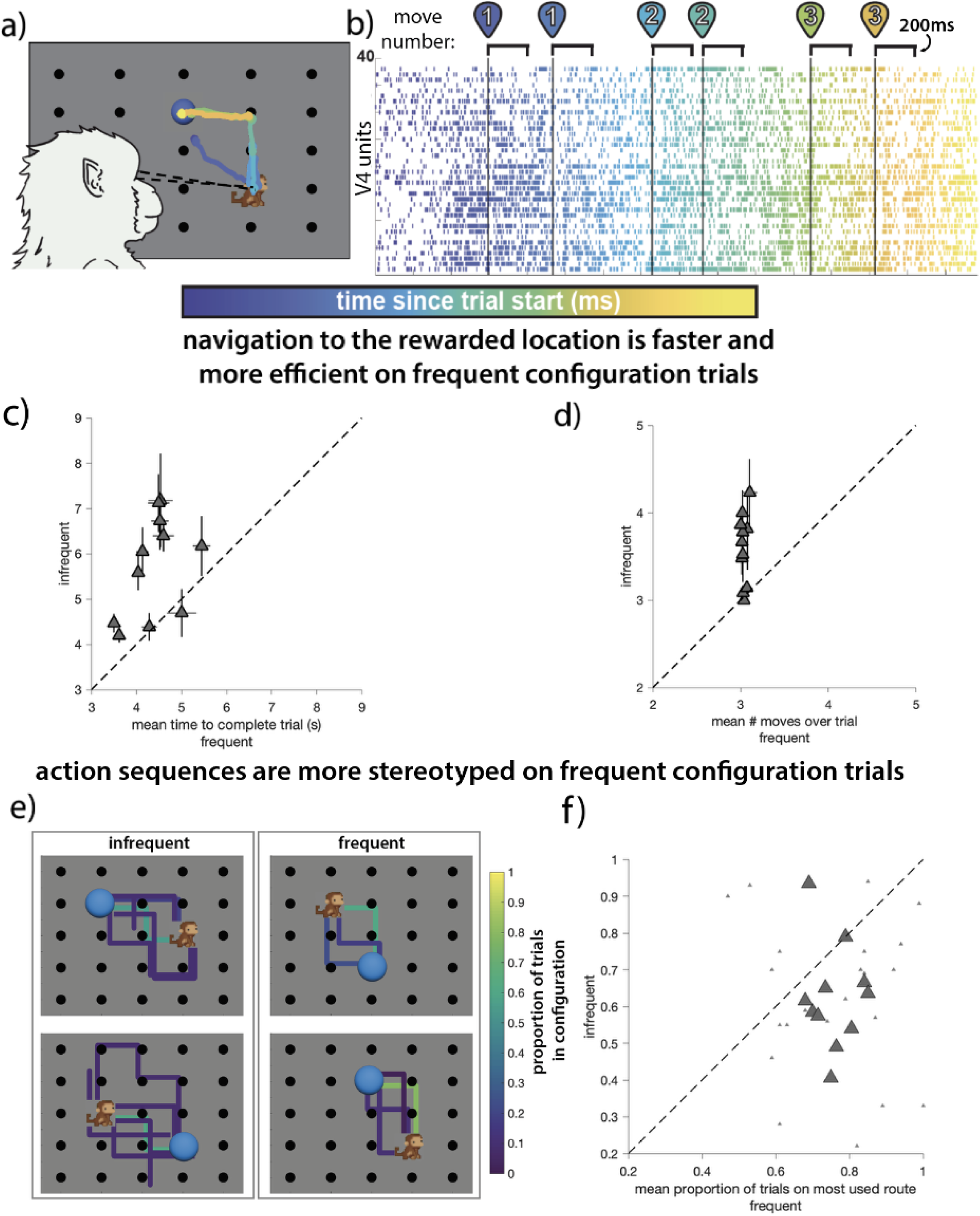
Active practice of action sequences. **a) Task design.** On each trial, the animal begins by fixating on a central point for 300ms. A grid of targets then appears and the animal uses eye movements to move the game piece (monkey icon) to the location of the reward (blue ball). To make a move, the animal must first fixate on the game piece for 200ms, then make a saccade to one of the four targets adjacent to the game piece. The trial ends with reward when the game piece reaches the target location. Trials terminate without reward if not completed within 20 s. **b) Electrophysiology.** While the animal performed the task, we simultaneously recorded populations of neurons from area V4. Each move (numbered) in a trial was associated with two periods of stable fixation (200ms, brackets) following fixation on the game piece and the chosen target. **c) Average number of moves per trial for each session.** Each point is the mean number of moves the animal took to reach the rewarded location on frequent (x-axis) and infrequent (y-axis) configuration trials. Error bars are the SEM. **d) Average time to complete a trial for each session.** Each point is the mean number of seconds the animal took to reach the rewarded location on frequent and infrequent configuration trials. Conventions as in (c). **e) Proportion of trials each route was taken for in a single example session.** Each line represents all routes taken from start to reward for every configuration. Line color indicates the proportion of trials in a configuration each route was used. **f) Average proportion of trials for most commonly used routes in each session.** Larger triangles are the mean proportion of trials for the most frequently used routes (greenest lines from (f)) across configurations on frequent (x-axis) and infrequent (y-axis) trials. Smaller triangles are proportions for all pairs of frequent and infrequent configurations.

This design provided well-defined periods of stable fixation following both game-piece and choice selection (Figure 4b), which was critical for interpreting activity in V4, a visual area responsive during fixation and eye movements^51–53^. All neuronal analyses were restricted to these fixation epochs. The spacing of the grid was chosen so that the population receptive fields of recorded units overlapped the currently fixated target. We did not constrain the number or identity of allowed moves, enabling us to measure experience-dependent emergence of subject-led action sequences.

Each session included four start–reward configurations. Two configurations were frequent (80% of trials) and reused across sessions; two were infrequent (20%) and novel within each session. This structure allowed us to compare behavior and neuronal activity during well-practiced and minimally practiced configurations.

### Repeated experience promotes efficient and stereotyped action sequences

A hallmark of learning in sequential decision tasks is the emergence of routines, in which a series of individual choices becomes organized into a repeatable sequence that can be executed with reduced deliberation^54–58^. Importantly, routine formation does not require that a particular sequence be uniquely optimal or explicitly reinforced. Instead, routines often emerge through repeated experience with similar task demands, reflecting a shift from flexible exploration toward consistent reuse of familiar action sequences.

We therefore hypothesized that, with experience, the animal would increasingly organize visually guided decisions into stereotyped action sequences for frequently encountered configurations. Consistent with this prediction, frequent configurations required fewer moves to reach the rewarded location than infrequent configurations (Figure 4c; one-sided paired t-test: p = 4.0×10^−4^; effect = 0.56). Frequent trials were also completed more quickly (Figure 4d; p = 6.2×10^−4^; effect = 1.3).

To assess stereotypy directly, we enumerated all unique routes taken within each session (Figure 4e) and measured the proportion of trials on which the most common route was used (Figure 4f). The animal relied more heavily on the dominant route during frequent configurations (one-sided paired t-test: p = 0.011; effect = 0.13), consistent with experience-dependent formation of action routines.

These results establish a behavioral context in which temporal structure is learned and expressed in measurable changes in behavior. This setup allows us to examine how experience with action sequences reshapes population activity in visual cortex, and whether the organization of these representations reflects task-relevant variables and learned sequence structure.

### Practice constrains population activity without reducing firing rates

We next examined how repeated practice reshaped V4 population activity. In contrast to passive familiarity paradigms, we did not observe a significant reduction in mean firing rate during execution of frequently practiced sequences. Mean population spike counts during game-piece fixations were comparable for frequent and infrequent trials (Figure 5a; two-sided two-sample t-test: p = 0.12). Thus, improved behavioral efficiency was not accompanied by global response suppression.

**Figure 5.**
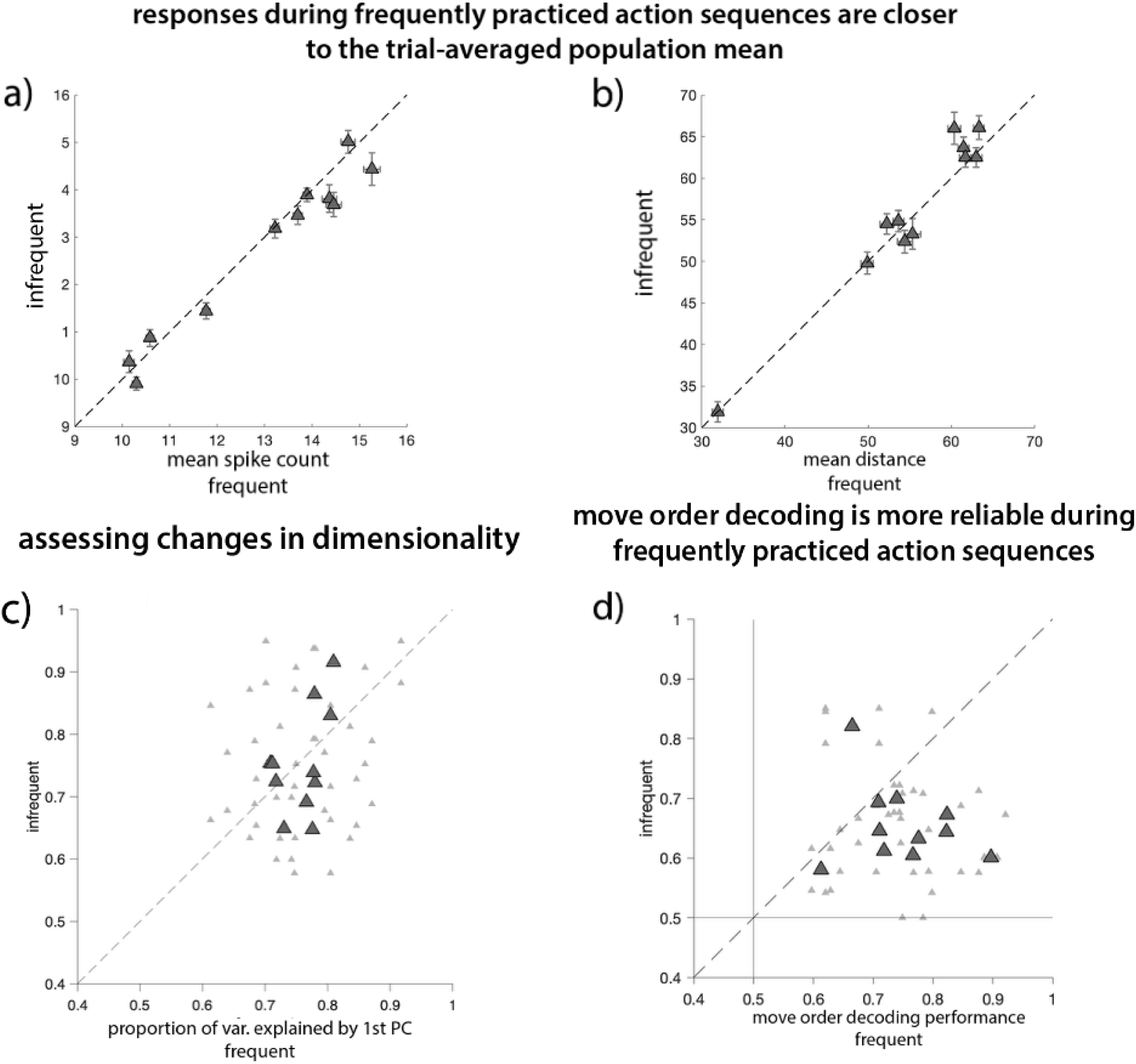
Active practice of action sequences reshapes population responses in visual cortex: **a) Average population spike count for each session.** Each point is the average spike count of the population over all fixations made on either frequent (x-axis) or infrequent (y-axis) configuration trials. Error bars represent the SEM. **b) Average Mahalanobis distance for each session.** Each point is the average distance from the trial-averaged mean population response to all fixations made on either frequent (x-axis) or infrequent (y-axis) configuration trials. Conventions as in (a). **c) Proportion of variance explained by the first dimension for each session.** Larger triangles are the mean proportion across both frequent or infrequent configurations. Smaller triangles are mean proportions for all pairwise combinations of frequent and infrequent configurations. Proportions found for frequent and infrequent choice fixations separately. **d) Move order decoding for each session using linear classifiers trained and tested on frequent or infrequent configuration trials.** Triangle size is the same convention as (c). Performance above 0.5 indicates above-chance decoding; values below the unity line indicate sessions where decoding was improved on frequent configuration trials.

However, practice was associated with a significant change in population geometry. Using Mahalanobis distance from the trial-averaged mean population response to quantify trial-by-trial dispersion, we found that responses during infrequent trials were significantly farther from the mean than those during frequent trials (Figure 5b; one-sided two-sample t-test: p = 0.022; effect = 1.1). This result indicates that practice constrains population activity even when overall response magnitude is preserved.

We then asked whether this constraint reflected a reduction in dimensionality. The proportion of variance explained by the first principal component did not differ between frequent and infrequent trials (Figure 5c; two-sided paired t-test: p = 0.78), indicating that experience-dependent constraint was not driven by collapse onto a lower-dimensional subspace.

Together, these findings reveal both similarity and divergence from the passive experiments. Practice constrained population responses toward a typical activity pattern, as measured by Mahalanobis distance, yet did so without reducing firing rates or dimensionality. Unlike the passive paradigms, however, this task provides a behavioral context in which sequence structure is directly relevant to performance. This allows us to investigate how experience-dependent reorganization of population geometry relates to the representation of behaviorally relevant variables.

### Practice strengthens representation of action sequence order

Although the task permitted flexible route selection, repeated experience led to stereotyped action routines. If the animal relies on such routines, maintaining information about position within the chosen sequence may be relevant. We therefore examined whether V4 population activity reflected relative move order.

Linear decoding analyses applied to choice-target fixations revealed that relative move order could be decoded significantly above chance for both frequent and infrequent trials (Figure 5d; frequent p = 6.2×10^−7^, effect = 0.25; infrequent p = 6.9×10^−7^, effect = 0.14). Temporal information about the action sequence was thus present in V4 regardless of familiarity.

However, decoding performance was significantly higher on frequent trials (Figure 5e; one-sided, paired t-test: p = 9.0×10^−4^; effect = 0.10), indicating that practice strengthened the linear representation of sequence position. This was not an artifact of simply having more trials in frequent than infrequent configurations; for this and all decoding analyses, we subsampled trials in frequent configurations to match the number of trials in the infrequent configurations. Practice therefore reorganized population geometry in a manner that enhanced expression of temporal order information without altering overall firing rates or dimensionality.

### Practice strengthens and differentiates representations of task-relevant variables

Repeated execution of action sequences constrained population activity in V4 and strengthened the representation of move order. We next examined whether this experience-dependent reorganization extended to additional task-relevant variables. Because successful performance depends on encoding reward location, distance to reward, and spatial position of the game piece, we asked whether practice altered how these variables were represented in population activity.

We compared linear decoding performance for four task variables during game-piece fixations on frequent and infrequent configuration trials. The variables were reward location (constant within a trial), distance to reward (updated after each move), and the X- and Y-position of the game piece within the grid. Across sessions and variables, decoding performance was significantly higher on frequent trials (Figure 6a; one-sided, paired t-test: p = 0.0013; effect = 0.073). Thus, repeated practice was associated with stronger linear representations of multiple task-relevant dimensions.

**Figure 6.**
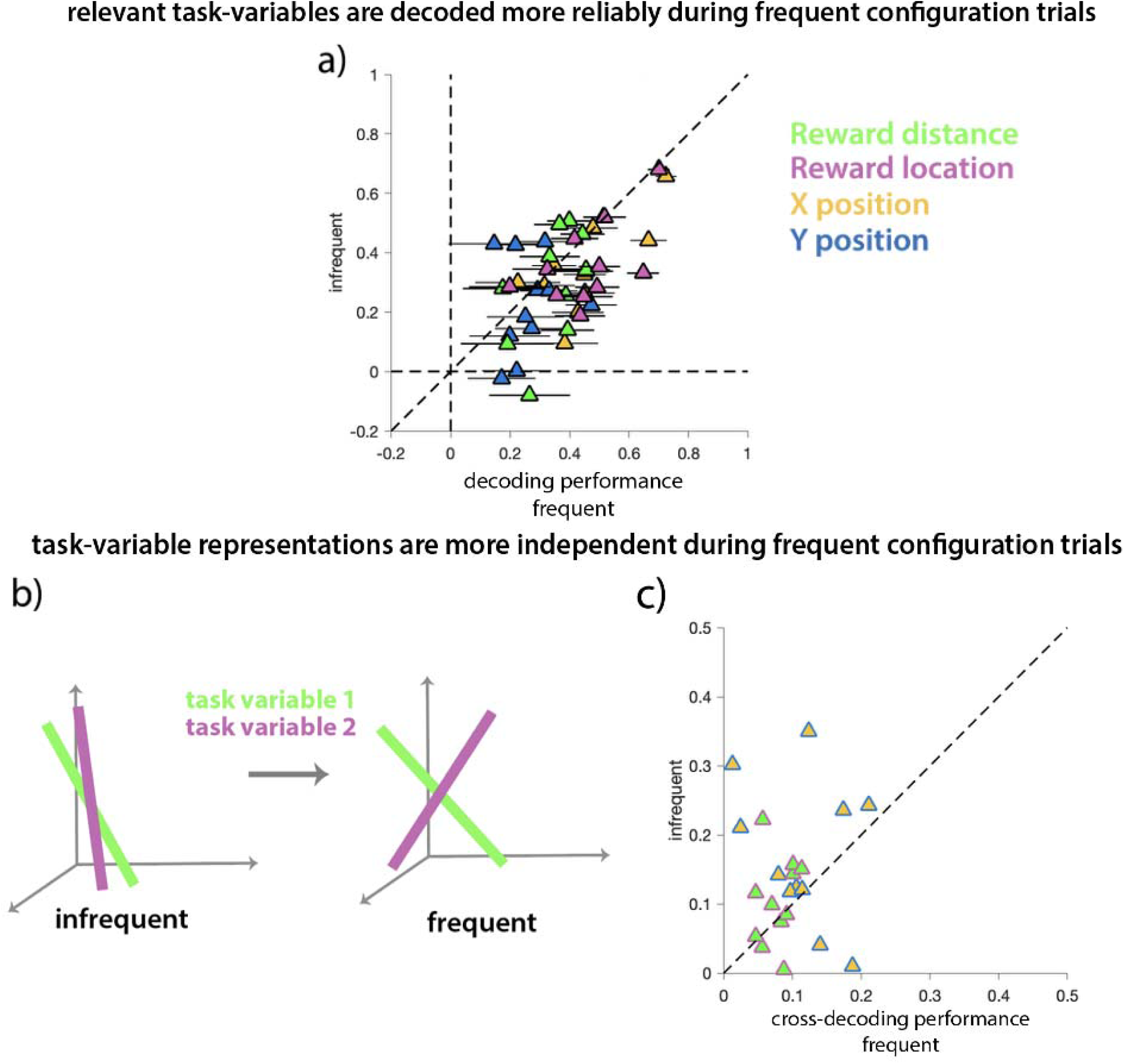
Active Action Sequence Practice: **a) Decoding performance of reward distance, reward location, game piece X-position and Y-position for each session.** Each point represents the correlation between actual and predicted values for each task-variable (colors) estimated using frequent (x-axis) or infrequent (y-axis) fixations. Horizontal error bars represent the SEM of the correlation coefficient taken across iterations of subsampled frequent fixations. **b) Schematic illustrating reduction in feature interference.** Uncorrelated task variables’ axes become more orthogonal, leading to more independent representations and a reduction in cross-decoding performance. **c) Cross-decoding performances for each session.** Each point represents the correlations between actual and predicted values for reward distance and Y-position, decoded using reward location and X-position axes, respectively. Colors represent pairings of features: gold and blue (X- and Y-position); green and magenta (reward distance and location)

We next examined whether practice altered the independence of these representations. Under conditions of high cognitive load, representations of task variables in V4 become less independent, which can lead to interference between variables and behavioral errors^25,26^. Based on this previous work, we hypothesized that repeated practice of action sequences might increase separability among task-relevant dimensions.

To test this hypothesis, we performed cross-decoding analyses in which a linear decoder trained on one variable was tested on another. Lower cross-decoding performance indicates greater independence between representations. Cross-decoding performance was significantly lower on frequent trials than on infrequent trials (Figure 6c; one-sided, paired t-test: p = 0.032). To ensure that this effect was not driven by poorly estimated decoding axes, we excluded sessions in which decoding performance (Figure 6a) for any variable in a cross-decoding pair was below 0.2. In this restricted subset, cross-decoding performance remained significantly lower on frequent trials (one-sided, paired t-test: p = 0.016; effect = 0.075).

Together, these results indicate that repeated practice reorganizes V4 population activity in a manner that increases linear separability among task-relevant variables. In contrast to passive familiarity effects, where experience was associated primarily with response suppression and constraint, practice in this behaviorally structured task was accompanied by selective strengthening and differentiation of task-relevant representations.

## Discussion

Across three forms of experience that differed in temporal structure, stimulus familiarity, and behavioral relevance, repeated experience was associated with systematic reorganization of population activity in visual area V4. In passive familiarity and sequence paradigms, experience was accompanied by reduced mean firing rates and decreased dispersion of population responses. In contrast, during active execution of practiced action routines, experience constrained population dispersion without reducing firing rates or dimensionality.

Despite these differences in response magnitude and representational dimensionality, all three contexts shared a common feature: more frequently experienced conditions were associated with population responses that lay closer to a typical activity pattern and, in structured tasks, more strongly reflected environmental regularities. These findings suggest that experience imposes a consistent organizing influence on visual population geometry, while also raising the possibility that the specific form of reorganization is tuned to the temporal and behavioral demands of the task.

### Experience constrains population activity in V4 within an asymmetric response space

Experience-dependent changes in response magnitude have been widely reported in sensory cortex, particularly in the context of novelty, surprise, release from adaptation, and violations of learned structure^1–4,7–14,43,44,59–62^. In our passive viewing experiments, V4 population responses were larger during first presentations of novel stimuli and when previously experienced image sequences were violated, consistent with previous work showing enhanced activity for unexpected stimuli or events.

However, neuronal firing-rate distributions are inherently asymmetric: responses are bounded below by zero but unbounded above. Responses that fall far from the typical population state are therefore more likely to do so via increases in firing rate. Under this view, novelty-related increases in activity may reflect population responses that deviate from a typical response distribution, rather than selective amplification of specific representations.

Multivariate measures that quantify the distance of population responses from their typical distribution provide a complementary perspective on these effects. Across all three experimental contexts, more frequently experienced conditions were associated with smaller Mahalanobis distance from the trial-averaged mean response. This pattern was observed whether mean firing rates decreased, increased, or remained unchanged. Thus, experience consistently constrained the dispersion of population responses within population space, even though the accompanying changes in firing rate differed across paradigms.

Importantly, this geometric account does not render firing-rate modulation irrelevant. Rather, it suggests that changes in response magnitude may often arise from shifts in how responses are distributed within a constrained response space. Quantifying population distance makes this relationship explicit by measuring how experience alters dispersion relative to the typical population state. In this way, population-level metrics offer a unifying framework for understanding why novelty and surprise are often associated with enhanced activity, while familiar conditions evoke responses that are both more typical and, on average, lower in magnitude.

### Experience embeds temporal structure in population geometry

Constraining responses toward a typical activity pattern is not equivalent to uniform compression. In contexts with temporal structure, experience was associated with systematic reorganization of population geometry that reflected sequence order.

In the passive sequence experiments, position within a learned image sequence became more linearly aligned in population space over the course of experience. In the action sequence task, move order could be decoded above chance in both frequent and infrequent conditions, but decoding performance was stronger for frequently practiced configurations. Thus, temporal information was present in V4 even under flexible choice, and experience strengthened its expression in population activity.

Notably, the form of reorganization differed between the passive and active contexts we studied. In passive sequences, experience was accompanied by reductions in firing rate and, in some analyses, dimensionality. In contrast, during active sequence execution, temporal information was strengthened without reductions in firing rate or dimensionality. These differences suggest that embedding temporal structure in population geometry does not require a single neural signature, and may instead reflect context-dependent adjustments in how population activity is organized.

Routine formation provides one possible interpretation of these effects. With repeated exposure, animals adopted stereotyped action sequences despite the availability of alternative routes. Reduced behavioral variability may stabilize transitions between successive states, allowing position within a chosen sequence to be represented more consistently. Under this view, experience-dependent constraint of population responses and strengthened temporal coding may arise together as aspects of a coordinated reorganization tuned to task demands.

### Representations of task-relevant information are strengthened and differentiated through practice

The action sequence task further allowed examination of how experience influences the representation of multiple task-relevant variables. In addition to move order, reward location, distance to reward, and spatial position of the game piece were linearly decodable from V4 activity. Decoding performance was consistently higher during frequently practiced configurations, indicating that experience altered the organization of population responses in a manner that strengthened linear representations of several variables.

Practice was also associated with greater separability among these representations. Previous work has shown that under task uncertainty or high cognitive load, representations of independent task variables in V4 become less separable, leading to interference between dimensions that should remain distinct^25,26^. Infrequent configurations in the present task can be viewed as relatively uncertain contexts. Consistent with this interpretation, cross-decoding performance between task variables was lower during frequent trials, indicating increased differentiation among representational dimensions with practice.

### Common principles across passive and active experience

Although the passive and action-based experiments differed substantially in behavioral relevance, task structure, and stimuli, they revealed some similarities in how experience reorganizes population activity. In all cases, experience constrained population responses while preserving or enhancing structure in how regularities in the environment are represented. In the passive sequence task, these regularities took the form of temporal order. In the action sequence task, they additionally reflected variables relevant to ongoing decisions and route planning.

This similarity across passive and active experience is consistent with a broad literature suggesting that a wide range of perceptual, cognitive, and motor processes have similar effects on the responses of neurons in visual cortex. Disparate processes including adaptation, contrast, attention, task switching, arousal, and saccade planning are all associated with the scaling of visual responses (gain changes^3,25,27,30–36,53,63,64^) and many are also associated with changes in the magnitude and structure of interactions between neurons^15,27,36,65^. The present findings extend this work by suggesting that such diverse effects may be understood within a common geometric framework: experience reshapes the distribution and organization of population responses, while the specific form of that reshaping may be tuned to the temporal structure and behavioral relevance of experience. Future work will be needed to determine how broadly these geometric principles generalize across tasks, brain areas, species, and timescales of learning, and to clarify how task demands shape the specific form of population reorganization.

### Future directions: experience-dependent organization of responses by ecologically-relevant categories

Our results suggest that experience can reorganize population responses to reflect temporal and task-relevant regularities. Similar principles may apply to other forms of environmental structure, including ecologically relevant categories. Stimulus distinctions such as animacy, real-world size, edibility, reachability, and conspecific identification are associated with systematic visual features and are known to influence cortical responses^66–76^.

If similar principles apply, experience may progressively organize population responses to reflect behaviorally meaningful structure across multiple domains. Understanding how population geometry evolves with experience may therefore offer a general framework for linking sensory representations to the temporal and behavioral structure of the environment.

## STAR Methods

### EXPERIMENTAL MODEL DETAILS

Five adult rhesus monkeys (Macaca mulatta; four males and one female; weights: M1 and M2 = 10 kg, M3 = 11 kg, M4 = 8 kg, M5 = 12 kg) were implanted with titanium headposts before any behavioral training. Because of the small number of subjects used in this experiment (typical of primate electrophysiology), we did not analyze differences between groups of animals. We therefore are not in a position to analyze sex as a biological variable.

### ARRAY IMPLANTATION

All animals were implanted with microelectrode arrays in cortical area V4, which was identified by visualizing sulci and using stereotactic coordinates. Experiments and surgeries for the passive image experience study (M1 and M2) were conducted at the Center for the Neural Basis of Cognition at the University of Pittsburgh/Carnegie Mellon University, and all animal procedures were approved by the Institutional Animal Care and Use Committees of the University of Pittsburgh and Carnegie Mellon University. Experiments and surgeries for the passive image sequence and active action sequence practice studies (M3, M4, M5) were conducted at the University of Chicago, and all animal procedures were approved by the Institutional Animal Care and Use Committees of the University of Chicago.

### PRE-ARRAY IMPLANTATION BEHAVIORAL TRAINING

Three of the animals participated in other experiments before being trained on the tasks presented here. However, the stimuli used in the current experiments had never been shown to any of the animals. Before array implantation surgery, all animals were familiar with the requirement to maintain fixation throughout a trial. M3 (active action sequence experiment) underwent a month-long period of training on a greatly reduced version of the task (two-by-two grid of stimuli) to learn the main goal of a trial (move the game piece to the reward location).

## METHODS DETAILS

### VISUAL STIMULI

In the passive image and image sequence experience tasks, grayscale images of animals and objects were sampled from a larger stimulus set used in Long et al., 2018. The passive image sequence and active action sequence experiments utilized uniquely colored stimuli varying in medial axis curvature and orientation that were sampled from a larger stimulus set used in previous experiments from our laboratory (ref, Ram). Visual stimuli were displayed using custom software (written in MATLAB using Psychophysics Toolbox (v.3)^77^) on 24” ViewPixx LCD monitors (1920 × 1080 pixels; 120 Hz refresh rate) placed 54 cm from each head-fixed animal. Eye position was recorded using an infrared eye tracker (EyeLink 1000; SR Research). We recorded neuronal responses and the signal from a photodiode (to align neuronal responses to the stimulus presentation period) using Ripple hardware.

### NEURAL RESPONSES

During all sessions, recorded electrical activity from each channel was filtered (bandpass 250-5000Hz) and threshold-crossing timestamps were saved. We also recorded raw electrical waveforms at each crossing. We use the term “unit” to refer to multiunit activity at each recording electrode. Analyses comparing single units and multiunits in our previous work^27^ as well as in another laboratory^78^, did not find systematic differences between single neurons and multiunits for the types of population analyses presented here.

In passive viewing experiments, stimulus-evoked V4 unit responses on each trial were calculated based on spike counts from stimulus onset through the duration of the stimulus presentation period. In the active action sequence experiment, stimulus-evoked spike counts were summed during stable periods of fixation beginning at the start of each game piece/choice fixation through the following 200 ms (before the next eye movement). For all experiments, baseline responses on each trial/fixation were calculated as the spike count during a 100–200 ms period of central fixation before stimulus onset.

### UNIT AND TRIAL/FIXATION EXCLUSION

For each recording session in all experiments, we excluded trials/fixations that were exceptionally noisy. To determine whether a fixation should be excluded from analysis, we computed the population response (mean spike count over units) for each fixation, z-scored those responses over all fixations, and excluded fixations for which the absolute value exceeded three standard deviations from the fixation-averaged population response. This fixation exclusion criterion allowed us to control movement artifacts.

We also excluded units that were exceptionally unresponsive. To determine whether a V4 unit should be excluded, we computed the mean (over all fixations) stimulus response (spike count during the first 100–200 ms, matched to baseline epoch length for each animal) and baseline response for each unit. We included units for which the mean stimulus response was at least 1.1× the mean baseline response. To exclude units with little to no recorded activity (during baseline or stimulus response), we calculated the total number of spikes for each unit over all fixations and excluded units with fewer than ten spikes over the entire session.

In one monkey, we observed instances of units becoming unresponsive over the course of a session. To determine whether this occurred, we computed the sum of spikes over all fixations in the first and second halves of each session for each unit and excluded units that showed 2.5× higher spike counts during the first half of the session than the second half.

The mean number of units included in each session for each experiment and monkey was: passive image experience data (M1 = 48 units; M2 = 40.3 units); passive image sequence data (M3 = 48.5 units; M4 = 63 units; M5 = 31 units); active action sequence data (M3 = 55.5).

### RECEPTIVE FIELD MAPPING

3D images of solid objects and Gabors were flashed in the appropriate hemifield of each animal following recovery from implantation surgery. A 5×5 grid of slightly overlapping locations was chosen, and two stimulus sizes were used to identify the location of each units’ receptive field. For each animal, a location within the joint receptive fields of the greatest number of units possible was identified for stimulus placement.

### EXPERIMENT ONE TASK DESIGN: PASSIVE IMAGE EXPERIENCE

On each trial, the animals were required to fixate centrally for 100–200 ms and maintain that fixation while the stimulus was presented for 860 ms. Following stimulus offset, the fixation spot disappeared and a target appeared at a radial location randomly sampled from a distribution of angles around the fixation point, but never overlapping the stimulus. The animal received a liquid reward for making a saccade to the target (eye movement provided as a control for low-level adaptation). One stimulus was presented per trial. Little training was required (< 1 week, if any) to achieve adequate task performance for both animals.

Each image was repeated in the trial following its first presentation. On average, the time between image repeats was 3.0 seconds (SEM 0.053) for M1, and 4.4 s (SEM 0.15) for M2. On any given session, stimuli were sampled from a larger stimulus set consisting of 120 images of animals and objects^70^. We analyzed sessions where at least 60 unique images were presented, resulting in five sessions for M1 and four sessions for M2.

### EXPERIMENT TWO TASK DESIGN: PASSIVE IMAGE SEQUENCE EXPERIENCE

#### Shortened sequence sessions

On each trial, the animals were required to fixate centrally for 100–200 ms and maintain that fixation while a sequence of three or four images was presented. Each image within a sequence was presented for 200 ms, with a 200 ms gap between images, resulting in a total stimulus presentation period of 1000–1400 ms. The animals received a liquid reward at the conclusion of each sequence.

Within a session, the expected sequence of four very familiar stimuli (top right, Figure 2a) was presented on 80% of trials. Randomly, on 20% of trials, a shortened, three-image version of that expected sequence was presented. In the unexpected sequence, the stimulus in the third position (C) was always omitted, resulting in the fourth-position stimulus (D) being displayed earlier than expected. We analyzed sessions in which at least ten unexpected sequences were viewed, resulting in four sessions from M3, three sessions from M4, and two sessions from M5.

#### Novel image sequence sessions

This experiment followed the same trial design as the shortened sequence sessions. However, on the random 20% of trials where there was a deviation from the expected sequence, one novel sequence of four images was presented per session. In sessions that utilized the grayscale images of animals and objects, the novel image sequence always followed the same categorical order as the expected image sequence (i.e.,big animal, big object, small animal, small object). As before, we analyzed sessions where at least ten unexpected sequences were viewed, resulting in five sessions from monkey M3.

### EXPERIMENT THREE TASK DESIGN: ACTIVE ACTION SEQUENCE PRACTICE

The animal learned that the goal of each trial was to move the game piece stimulus (monkey, Figure 4a) to the location of the reward stimulus (blue ball) before the trial terminated (20 seconds). To initiate a trial, the animal fixated on a central spot for 300–375 ms. A 5×5 grid of targets populated with the game piece and the reward stimuli was then displayed, the trial timer was started, and the animal was released from fixation requirements.

To make a move, the animal was required to fixate (for 200 ms) on the target associated with the game piece, make a saccade to one of the directly adjacent (up, down, left, or right; not diagonal) targets, and fixate on that chosen target (for 200 ms). If the animal executed a move correctly, the position of the game piece would update to the chosen location. At the start of every trial, the game piece and the reward were always separated by three moves. The animal could make as many moves as desired, and a juice reward would be dispensed as long as the game piece reached the location of the blue ball before the trial terminated.

Within a session, on 80% of trials, game piece start and reward locations were displayed in one of two extremely familiar configurations. These frequent start and reward configurations were shown to the animal on the majority of trials across all sessions. Randomly, on 20% of trials, one of two other possible start and reward location configurations was shown. Both infrequently shown configurations were completely novel to the animal each session. We analyzed sessions in which at least ten trials of each infrequent stimulus configuration were completed. Eleven sessions from M3 passed this criterion.

### QUANTIFICATION AND STATISTICAL ANALYSIS

In the passive experiments, we quantified responses for each unit by summing the number of spikes over the stimulus presentation period for every trial (one stimulus/trial in experiment one, three or four/trial in experiment two). In the active action sequence practice experiment, spike counts were summed across the 200ms following each fixation on the game piece or choice target. Unless otherwise stated, all statistical tests were paired.

#### Finding pairs of D/D’ presentations (Figure 2)

To more accurately compare responses to presentations of expected (D) and unexpected (D’) positions of the last stimulus in the shortened sequence sessions, we paired presentations that were closest together in terms of session time. For each unexpected presentation of D’ in a session, we found the closest preceding instance of D.

#### Calculating mean population spike counts (Figures 1b-c, 2b-c, 5a)

In all experiments, we calculated the population spike count for each stimulus presentation or fixation by finding the average response across all V4 units (Figures 1b, 2b). For each session, we then took the mean of those population responses over all trials/fixations in a condition (i.e., first/second, D’/D, infrequent/frequent; Figures 1c, 2c, 5a). For statistical analysis of the active action sequence experiment, we performed two-sample tests on pooled (across sessions) population responses to infrequent and frequent fixations.

#### Calculating Mahalanobis distance (Figures 1e, 2d-e, 5b)

In all experiments, we calculated mahalanobis distances from a matrix of z-scored responses for all units and trials/fixations [all trials/fixations x all units] (Figure 2d). We found the mean distance over all trials/fixations in each condition for each session (Figures 1e, 2e, 5b). For statistical analysis of the active action sequence experiment, we performed two-sample tests on pooled (across sessions) distances for infrequent and frequent fixations.

#### Calculating sequence population responses (Figure 3a)

For all expected trials in all passive image sequence sessions (shortened and novel violations), we found the sequence population responses by averaging over population spike counts to all images in the sequence. We then found separate averages of those sequence population responses over the first and last twenty five trials in each session.

#### Calculating the sum of residuals and slopes for linear fits to image sequences (Figures 3c-d)

For all passive image sequence sessions we found the mean population response to each image in the expected sequence over all trials, and found the line-of-best-fit for these four mean responses. On every trial, we subtracted that trial-averaged image response from the population response to each image, and took the sum of the residuals (sum of squared errors) to that line of best-fit for all images. This procedure left us with one sum of residuals for each trial. We then fit another line to those residual values over the course of a session (Figure 3c), and calculated the slope. We plotted the distribution of those slopes for every session, and took the mean across sessions (Figure 3d).

#### Estimating dimensionality of population responses in each experience condition (Figures 3e, 5c)

For all passive image sequence (expected trials only) and active action sequence sessions, we first split up trials/fixations by experience condition (early/late, infrequent/frequent). We then separately calculated the proportion of variance explained by the first principal component for trials in each condition.

#### Decoding temporal order information (Figures 3f, 5d)

##### Passive image sequence sessions (Figure 3f)

For all expected trials in all passive image sequence sessions, we performed separate decoding analyses to compare sequence order readout during the first and next twenty-five trials within each session (early vs late experience). First, we separated responses to all sequence images (A, B, C, D) into early and late trials. Within each experience condition, responses were z-scored across trials for each unit.

We then constructed three complementary train–test splits designed to assess whether sequence elements were arranged along a consistent ordering axis in population space: (A+B vs C+D), (B+C vs A+D), and (C+D vs A+B). For each split, we trained a **linear classifier** to distinguish the two groups of sequence elements using cross-validation within the experience condition. The learned decision boundary was then tested for its ability to generalize to classify the complementary grouping of elements in a manner consistent with sequence order.

Performance was scored in a sign-invariant manner, such that correct ordering of elements (A, B, C, D) or its reverse (D, C, B, A) were treated equivalently. This procedure tested consistency of monotonic ordering rather than absolute temporal labels. Chance decoding performance was 0.5.

For each session and experience condition, we averaged decoding performance across the three splits and compared early and late trials (Figure 3f).

##### Active sequence practice sessions (Figure 5d)

For all active action sequence practice sessions, we performed separate decoding analyses to compare relative move order readout during infrequent and frequent trials. Analyses were limited to the first three fixations within each trial because three moves was the minimum number required on all trials. Here, A, B, and C refer to the first, second, and third move positions, respectively.

First, we separated trials by unique start and reward location configuration (four configurations per session: two frequent and two infrequent) and z-scored responses within each experience condition. Within each condition, we trained and tested linear classifiers using cross-validation.

To assess generalization across move positions, we constructed two train–test splits: (train on responses distinguishing A vs B and test on C) and (train on B vs C and test on A). These splits evaluated whether the learned decision boundary generalized consistently across move positions, consistent with a monotonic representation of move order.

As in the passive analysis, decoding performance was scored in a sign-invariant manner to test consistency of ordering rather than absolute labels. Chance performance was 0.5. We averaged decoding performance across splits and across start/reward configurations within each experience condition and compared infrequent and frequent trials (Figure 5d).

#### Task-variable decoding in active action sequence experiment (Figure 6a)

In each session, we isolated fixations from infrequent and frequent trials. We found reward distance values for each fixation by calculating the minimum number of moves that separated the game piece from the reward. For infrequent trial fixations, we then found the decoding performance for all four task-variables using linear regression. We cross-validated our model by leaving out one fixation to test and training on all others in the experience condition. We then calculated the correlation between predicted and actual task-variable values across all fixations. To match trial counts used for regression across the experience conditions, we randomly subsampled groups of frequent trials and performed the same cross-validated regression procedure 100 times.

#### Cross-decoding in active action sequence experiment (Figure 6c)

We performed cross-decoding for pairs of task-variables that were on average, uncorrelated across sessions (r<0.2). We used the same cross-validated linear regression procedure described above to predict reward distance values from the reward location axis for held-out fixations from infrequent and frequent trials. We repeated this process to predict Y positions from the X position axis.

## Supporting information

Supplemental Figures and Descriptions

## DATA AND CODE AVAILABILITY

Raw data is available at https://doi.org/10.6084/m9.figshare.31721140 and original code is available at https://github.com/lily-kramer/sequential_experience_reshapes_visual_reps.

## Acknowledgments

We thank Douglas Ruff, Cheng Xue, Ram Srinath, Grace DiRisio and Drew Sheets for comments on early versions of the manuscript and suggestions for data analysis. We thank the Kursti Eaves and the animal care staff at all institutions for help with the animals. This work was supported by the Simons Collaboration on the Global Brain award 542961SP1, NIH awards R01EY022930, R01EY034723, and RF1NS121913 to M.R.C.

