## Supplemental Figures and Descriptions for "Sequential experience reshapes population representations in visual cortex"

*Passive Image Experience Experiment*

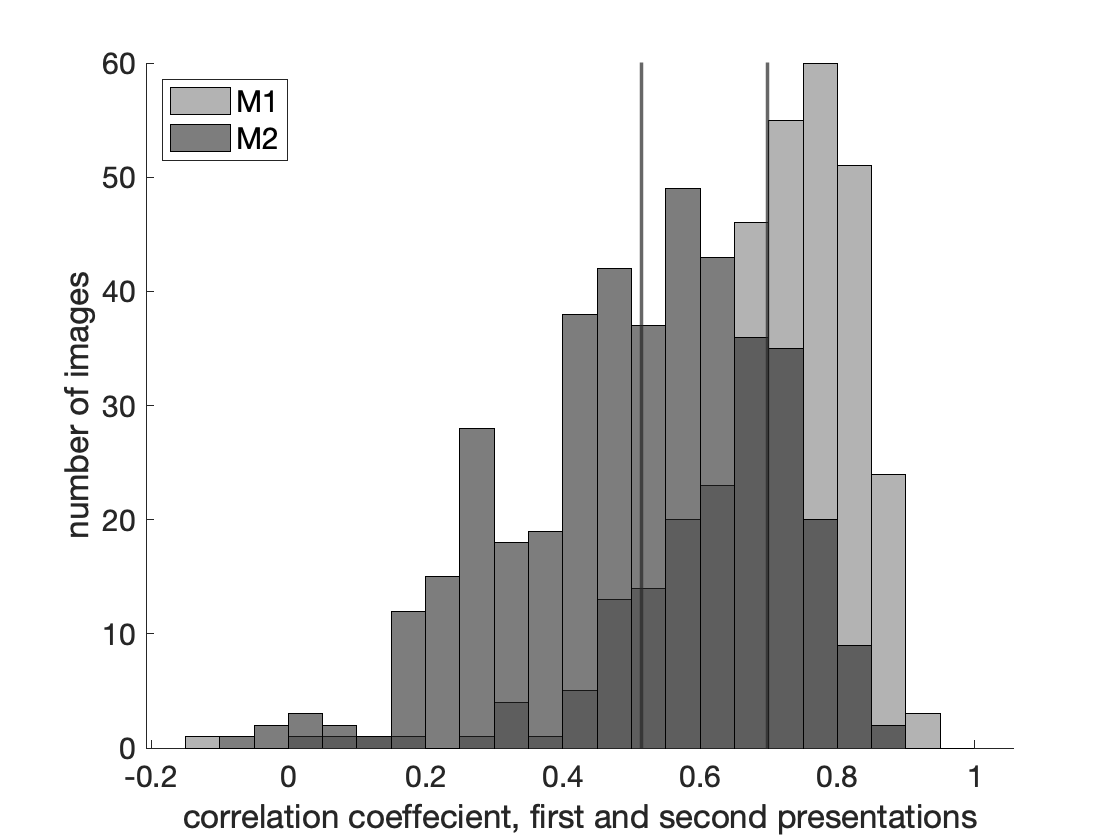

**Supplemental Figure 1: Correlations in population responses to first and second image presentations.** For all images, we found the correlation between population responses to first and second presentations. Means of distributions for each animal are represented by vertical lines (M1 mean = 0.70, SEM = 0.0081; M2 mean = 0.51, SEM = 0.0089).

*Active Action Sequence Experiment*

To control for differences in orientation of configurations (e.g., if infrequent configurations were primarily horizontal in comparison to the vertical frequent configurations), we limited our analyses to a subset of seven sessions where there was at least one vertical infrequent configuration tested. We redid analyses on this limited subset, and found that our results are in fact attributable to experience rather than configuration orientation, as evident by all metrics trending in previously reported directions (number of moves p = 0.0021; time to complete p = 0.0043; Mahalanobis distance p = 0.083; task-variable decoding p = 0.014; cross decoding p = 0.046).

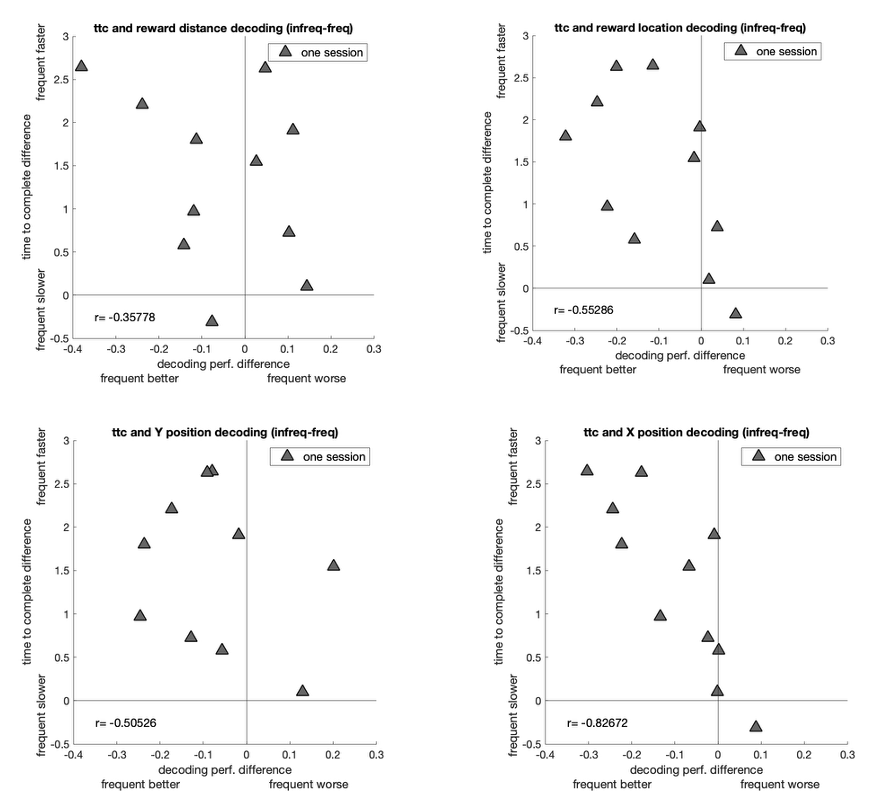

**Supplemental Figure 2: Correlation between differences time to complete and decoding performance across conditions.** For each session, we found the difference in mean time to complete and task-variable decoding performances across configurations (infrequent – frequent). A negative correlation indicates that for sessions where frequent trial completion times are much faster, decoding performance is also relatively better.
